# Characterizing drug resistance using geometric ensembles from HIV protease dynamics

**DOI:** 10.1101/379958

**Authors:** Olivier Sheik Amamuddy, Nigel T. Bishop, Özlem Tastan Bishop

**Affiliations:** Research Unit in Bioinformatics (RUBi), Department of Biochemistry and Microbiology, Rhodes University, Grahamstown 6140, South Africa; Department of Mathematics (Pure & Applied), Rhodes University, Grahamstown 6140, South Africa

## Abstract

The use of antiretrovirals (ARVs) has drastically improved the life quality and expectancy of HIV patients since their introduction in health care. Several millions are still afflicted worldwide by HIV and ARV resistance is a constant concern for both healthcare practitioners and patients, as while treatment options are finite, the virus constantly adapts and selects for resistant viral strains under the pressure of drug treatment. The HIV protease is a crucial enzyme that processes viral polyproteins into their functional form, and has been a game changing drug target since the first application. Due to similarities in protease inhibitor designs, drug cross-resistance is not uncommon across ARVs of the same class. It is known that resistance against protease inhibitors is associated with a wider active site, but results from our large scale molecular dynamics analysis further show, for the first time, that there are regions of local expansions and compactions associated with high levels of resistance conserved across eight different protease inhibitors visible in their complexed form in closed receptor conformations. The method developed here is novel, supplementary to the methods of nonsynonymous mutation analysis, and should be applicable in analyzing the structural consequences of mutations in other contexts.

## Introduction

Antiretroviral (ARV) drug resistance still persists despite recent improvements in antiretroviral therapy^1^. As the viral genome continues to accumulate mutations under the selective pressures of therapy^2^, surviving viral populations inevitably become less sensitive to one or more drugs over time. HIV reservoirs and its existence as a quasispecies^3^ means that an ARV should ideally inhibit a pool of slightly different conformations of receptor targets. The decreasing efficacy of drug binding over time means that patients may have to switch to more difficult treatment regimens with the possibility of experiencing more severe side-effects if no better-tolerated alternative exists. At the same time, ARVs are a finite resource which should be used with proper timing failing which resistance develops sooner. In order to design more robust ARVs and/or improve onto existing resistance prediction methods, additional knowledge of the motions associated with resistance may be helpful. However this is not straight-forward - patterns of resistance mutations patterns in HIV are complex^4^, involving mutations close to or away from the active site.

In this manuscript, we focus on HIV protease, which is a crucial enzyme for viral maturation, and is a well-established HIV drug target^5,6^. There are minor differences as to how various functional segments of HIV protease are defined in literature, possibly due to its high variability. Some of these structural features are annotated in Fig.1. Protease inhibitors (PIs) are competitive inhibitors of the enzyme^7^, which under normal circumstances processes the viral polyproteins Gag and Gag-Pol^8^. Multi-drug resistance within the PI class is not uncommon due to their long period of use and their three-dimensional and electrostatic similarities^9,10^. Previous work describe distinct mechanisms associated with ARV drug resistance that all point to active site expansion, namely (1) impaired hydrophobic sliding shown in the G48T/L89M double mutant with saquinavir^11^, (2) reduced dimer stability in L24I, I50V and F53L mutants^12^ and (3) single or co-operative distal mutations^13,14^.

**Figure 1.**
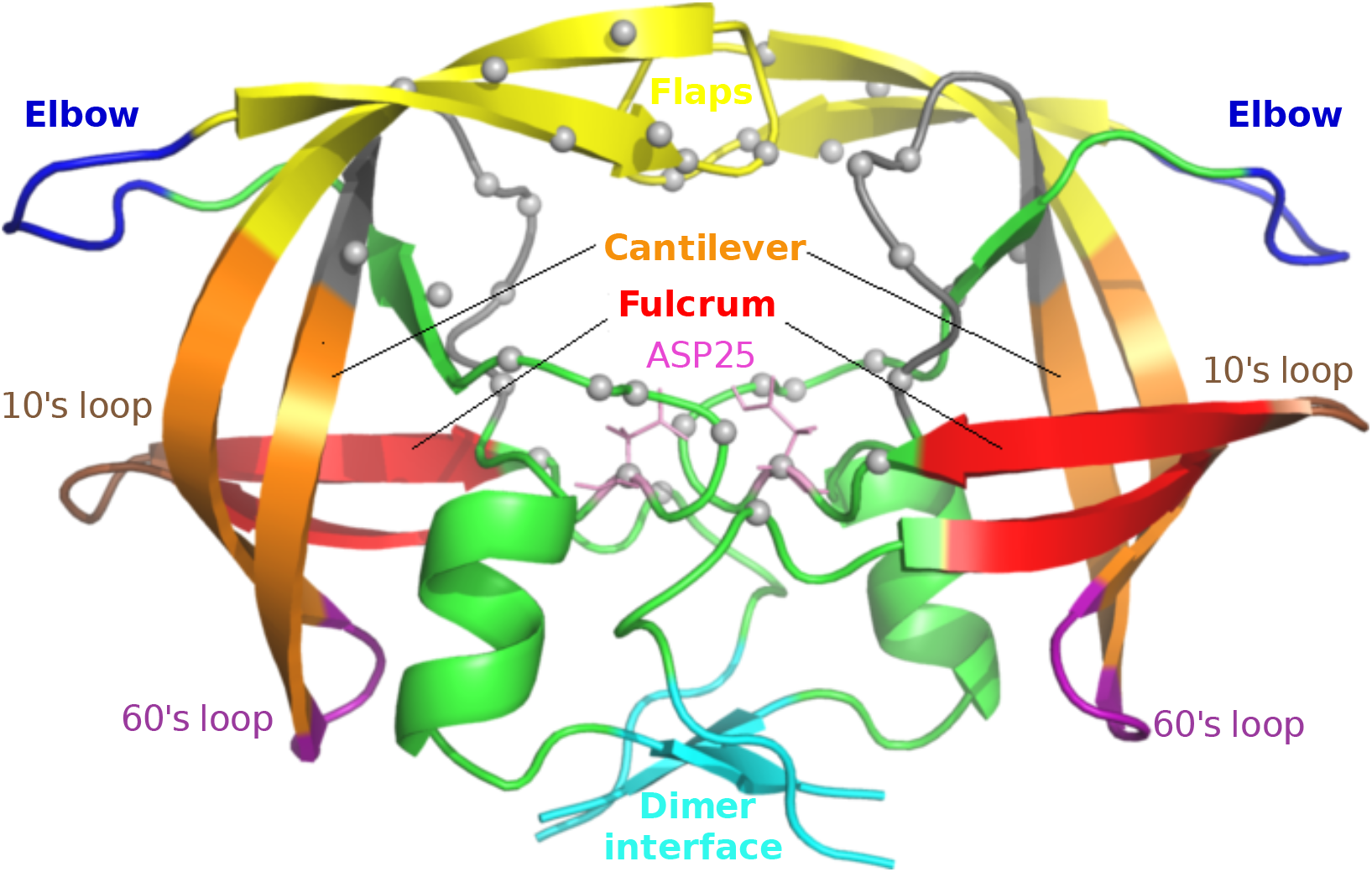
Functional regions within HIV protease. Grey spheres depict the residues constituting the binding cavity.

In resource-available settings drug efficacy can be inferred by monitoring viral load or CD4 cell counts, yet when available, knowledge of genotypic information can improve the choice of therapy to be used^15^. However, drug resistance mechanisms in HIV are not fully-understood^16^. Previous research has evaluated various computational modelling approaches over the years in order to predict or understand drug resistance mechanisms in HIV protease, ranging from the use of triangulations from static protease structures^17^, molecular docking^18,19^ to elastic network modelling^20^, molecular dynamics (MD)^19,21^ and many more, as reviewed by Cao and co-workers^22^. Here we adopt a structural approach using MD applied to 100 highly-resistant and 100 hyper-susceptible HIV sequences against eight protease inhibitors using the raw data available from the Stanford HIVdb^23^. We focus on the majority subtype (B), with the sequences containing rare residues removed, as described in our previous work^24^. 3D structures of the 200 protease sequences are built using homology modelling^25^ with a common drug-bound template for each target. Ligand docking with the eight ARVs, namely atazanavir (ATV), darunavir (DRV), fosamprenavir (FPV), indinavir (IDV), lopinavir (LPV), nelfinavir (NFV), saquinavir (SQV) and tipranavir (TPV) then gives 1,600 3D structures of drug-bound protease complexes.

Proteins are in constant motion^26^ and drug-binding alters their dynamics^27^ so each case requires its own MD run. Allowing for replications, a total of 3,200 MD runs are performed. Each run is about 2ns, so that in total the MD simulations amount to about 6,400ns. The observation of conserved resistance-related dynamics across this high number of independent short simulations of PI-bound receptor complexes shows that highly drug-resistant sequences can have structurally-detectable features. Considerable amounts of conformational sampling are typically required to observe motions that are of large amplitude^28,29^ or rare^30^. Same applies for increasing the accuracy of binding free-energy estimations (for instance between a ligand and a receptor), which comes with increased computational costs^31,32^. We circumvent these issues in this context, by describing two short and specific motions that are detectable very early in all-atom dynamics simulations of the retroviral protease. This would be very helpful if similar motions were to be picked up in non-B subtypes of varying residue composition as our algorithm highlights the most definitively distinct motions between drug resistance and susceptibility.

The *in silico* methods used are partially stochastic^33–35^, and we mitigate chance events by calculating statistical properties of each ensemble and applying network centrality measures. Remarkably, we find structural features of HIV protease that are different in the susceptible and resistant sequences, with the differences conserved across all eight ARVs. The results can form the basis for more robust ARV design and better prediction of drug resistance. Further, the combination of molecular dynamics, network centrality and statistical analyses used here provides an alternative way of analyzing the effects of non-synonymous mutations, and should be applicable to other diseases.

## Results and Discussion

As a quality control for all the MD simulations, *C_α_* RMSD values were first computed to exclude any error in periodic boundary corrections. A condensed representation of the mean and the standard deviations of RMSD values for each ARV is depicted in *SI Appendix*, Fig. S1. The runs were found to display slightly higher variation (in red) for the first 100ps before stabilizing (yellow to white) thereafter in each case. We then begin the experiment with a global assessment of the distributions of protein compactness using the radius of gyration (R_g_) across drug ensembles, as shown in Fig.2 to more local evaluations, namely pairwise residue distances and *C_α_* angles from receptors (Fig.3 to 6).

**Figure 2.**
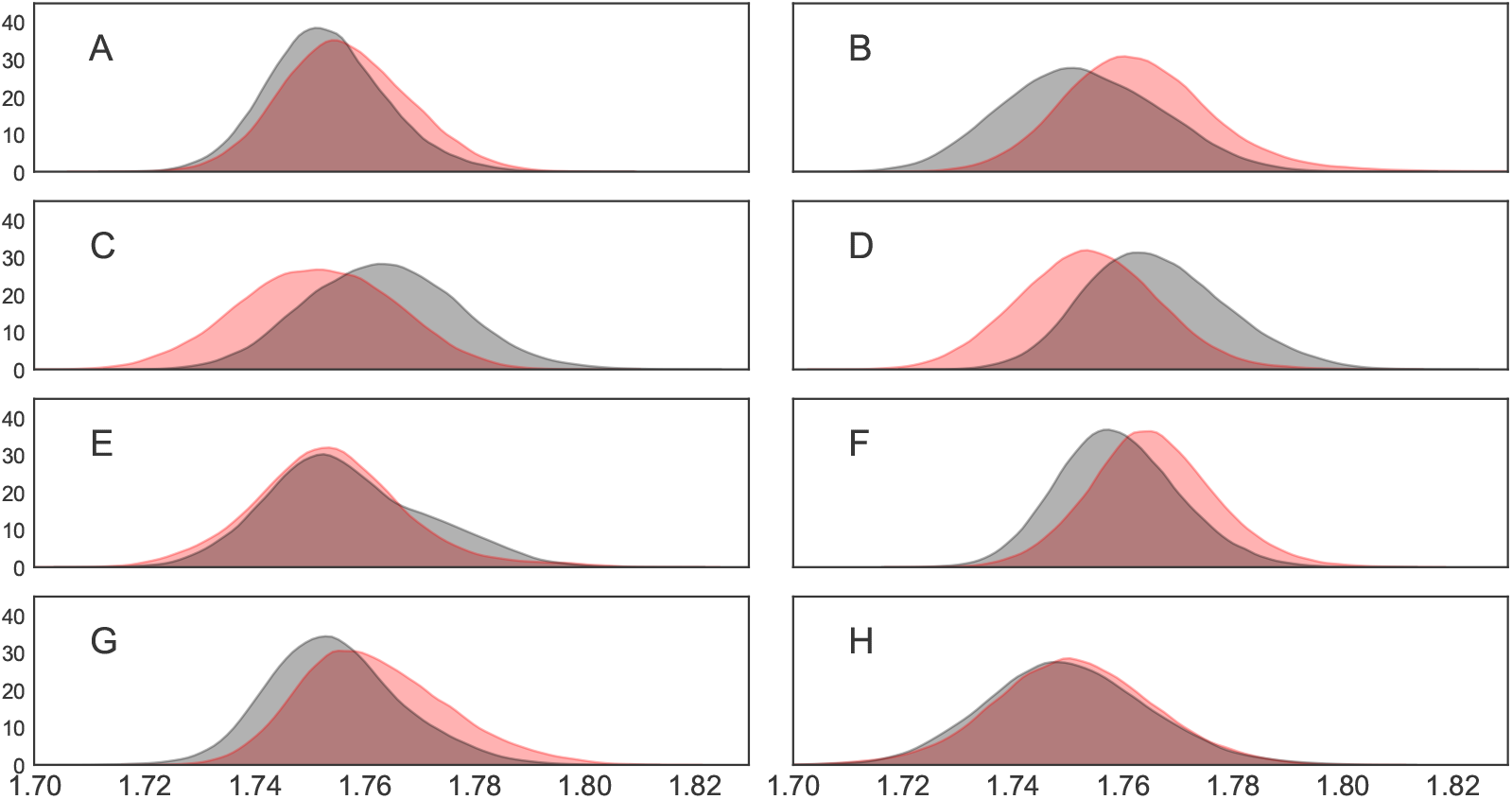
Distributions of *R_g_* values for protease inihibitor complexes containing ATV (A), DRV (B), FPV (C), IDV (D), LPV (E), NFV (F), SQV (G) and TPV (H). Resistant ensembles are shaded in red while susceptible ensembles are in grey.

**Figure 3.**
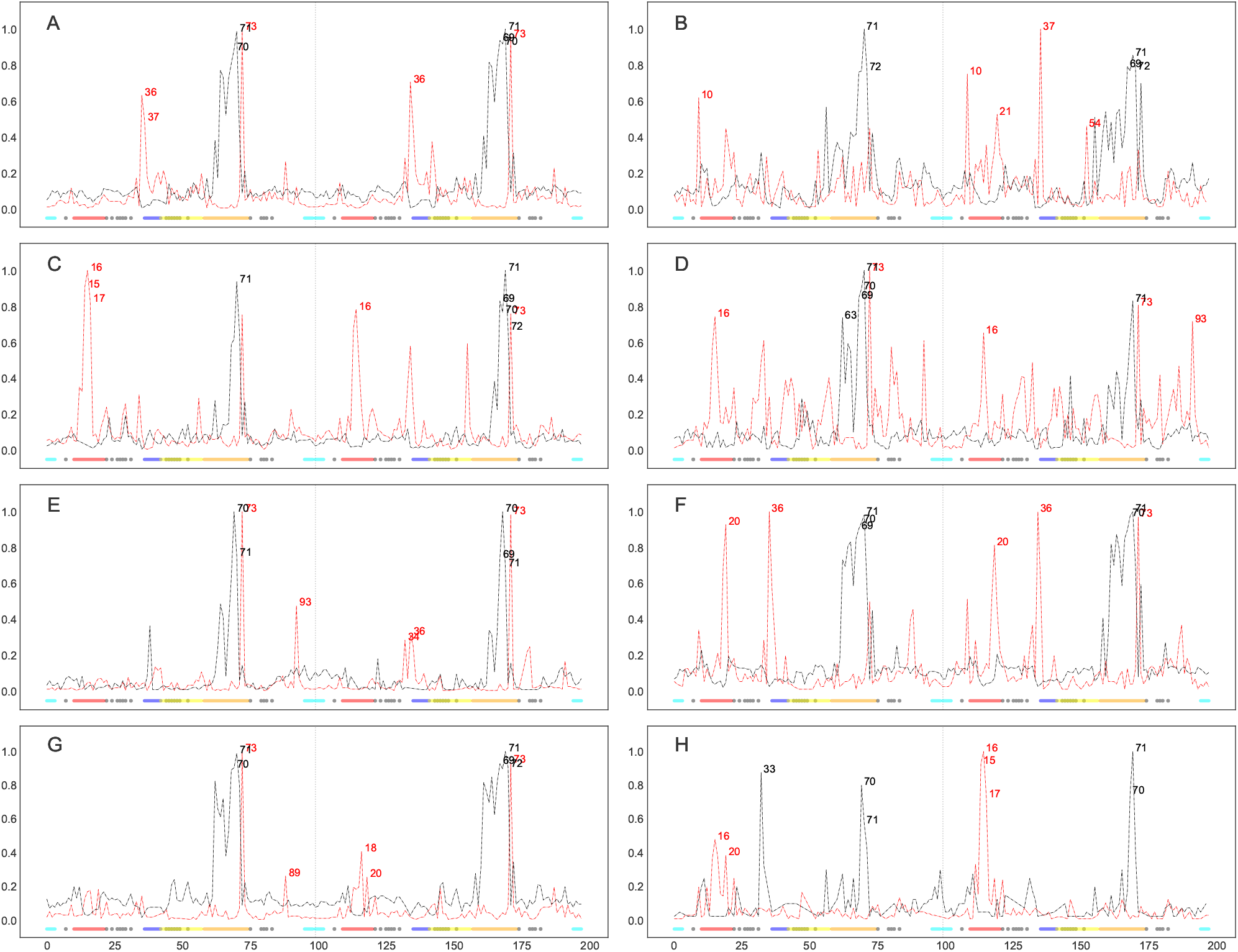
Normalized degree centralities of significantly larger (red lines) and smaller (**black lines**) distances observed in resistant ensembles for 8 FDA-approved protease inhibitor complexes, namely ATV (**A**), DRV (**B**), FPV (**C**), IDV (**D**), LPV (**E**), NFV (**F**), SQV (**G**) and TPV (**H**). The top 5 residue positions with the highest connectivities are labelled at the peaks in each graph. Inserted underneath are the functional protease residues depicted as colored dots, namely the fulcrum (red), the elbow (blue), the flap (yellow), the cantilever (orange), the interface (cyan) and the binding cavity residues (grey).

Contrary to what would be expected, the simulations showed no global tendency towards a less compact (larger *R_g_*) series of conformations in the resistant ensemble compared to the drug-susceptible state, as shown in Fig.2. Taken individually, only DRV, NFV and SQV display larger R_g_ values in their resistant ensemble. The reverse is actually observed in FPV and IDV. No appreciable shifts were observed for ATV, LPV and even less for TPV. Of notable interest are the slight shifts in the means of the R_g_ distributions across all drug ensembles, which are suspected to either be inherited from the template used for modelling or the docked drugs themselves, which could be propagating a different set of local receptor-ligand signals towards various parts of the receptor. Similar trends were observed upon replication (*SI Appendix*, Fig. S2). We hypothesize that compactions may instead be observable at a residue level and that these might be masked by more chaotic motions. Therefore we further investigate each protein-ligand complex by performing pairwise t-tests from residue distances obtained from aggregations of independent MD simulations.

The results of the Bonferonni-moderated t-tests are transformed into a network graph as explained in the Materials and Methods section, and represented as normalized degree centrality plots onto which several architectural features of HIV protease are mapped using the coloring scheme from Fig.1. Time averages of the distances for each residue pair are combined across proteins within an ensemble, instead of using the actual distances, mainly for computational efficiency. For all MD simulations, an initial region of higher RMSD fluctuation (100 ps) was discarded to reduce residual effects coming from prior equilibration. We further filter out stochastic variations by basing ourselves on the network concept of preferential attachment, which is the tendency of scale-free networks to attach new nodes to those that already have high connectivities^36^. The five top-ranked residue positions are subsequently prioritized on the basis of their highest connectivities for their higher support of being further away or closer with respect to a larger number of residues showing statistically-significant larger or smaller distances across the ensembles.

It is remarkable to note that despite the stochasticity of different sections of the experiment compounded with protein variations, our results show features common to all drug complexes. The base of the cantilever (very close to the 60’s loop) is drawn closer to the catalytic core for all drug complexes (Figs.3 and 4) in the resistant state. All the mutations present in each ARV’s resistance ensemble are shown in *SI Appendix*, Table S3, where known accessory and major drug resistance mutations (DRMs)^37^ are shown in bold black and red fonts respectively. From our simulations, a similarly conserved behavior is not immediately apparent from the degree centralities of those residues with larger distances in the susceptible ensemble, but can be seen from their mapping onto protease 3D structures. A lateral widening involving the elbow region and/ or the 10’s loop of the fulcrum is generally observed across all PIs. Part of this motion is described by Hornak and co-workers^38^ as events leading to flap opening, involving a concerted downward motion of the cantilever, fulcrum and flap elbow with an upward motion of the catalytic aspartate from the floor of the binding cavity. Further, we observed that residues of the receptor cavity do not appear amongst any of the top-ranked residues for each drug and ensemble, and behave in a quite opposite manner, with very low degree centralities. Occasional spikes did manifest themselves for some cavity residues, but these can be ignored as they may be chance events that would not be ranked similarly upon replication of the experiment, as seen in *SI Appendix*, Fig. S4. Low degree centralities in both ensembles (i.e. neither larger in the resistant nor in the susceptible ensembles) would point to the fact there is no consistent motion within the binding cavity that would define the state of drug-resistance or susceptibility, at least not within the time limit and conformational landscapes explored. This hints at a receptor pocket that is very malleable with multiple internal cavity dynamics that can lead to similar states, both within and between PI drug classes. A second scenario that could result in such low connectivities would be that cavity residues move in a coordinated manner across ensembles irrespective of drug exposure, which is unlikely. We now describe some ARV-specific results.

**Figure 4.**
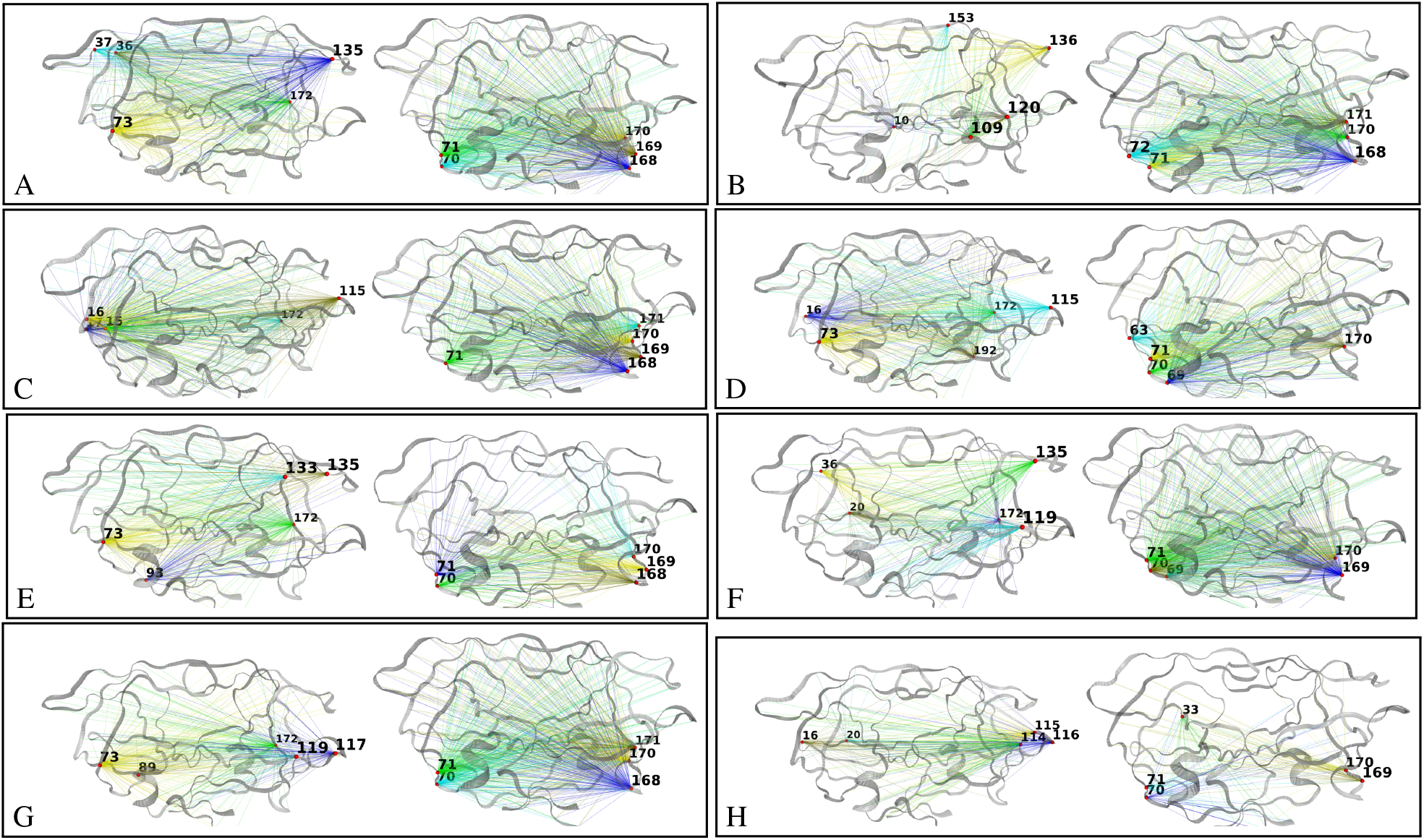
Mapping of the edges for top-ranked degree centralities onto HIV 3D protease structures for the significantly larger (left) and smaller (right) distances observed in resistant ensembles for 8 FDA-approved protease inhibitor complexes, namely ATV (**A**), DRV (**B**), FPV (**C**), IDV (**D**), LPV (**E**), NFV (**F**), SQV (**G**) and TPV (**H**).

In Fig.3A for ATV residues showing smaller distances in the resistant ensemble include positions 70, 71 (on chains A and B) and position 69-71 (chain B), while larger distances are at positions 36, 37, 73 (chain A) and at positions 36, 73 (chain B). Mapping these positions onto protein structures (Fig.4A) shows that regions predicted to be larger in the resistant ensemble move in a lateral outward direction, favoring a wider conformation, while regions predicted to be smaller in the resistance ensemble (or larger in the susceptible ensemble) show an upward motion with respect to the flaps. The residues involved in widening and shortening around the binding cavity show a high level of symmetry between each monomer of the protein. A very similar profile was obtained upon replication, with residue 36 peaking from within chain A instead of chain B. Such conservation in behavior may find direct application in drug resistance prediction or for feature augmentation for improving machine learning prediction of resistance. Mutations present in the ATV resistance ensemble include both accessory DRMs 10IVF, 32I, 33FV, 34Q, 46LI, 48V, 53L, 54LVM, 60E, 62V, 64VM, 71IV, 73STA, 90M, 93LM and a major DRM 84V, in addition to multiple other variations.

In the case of DRV (Figs.3B and 4B), residues at positions 71 and 72 (chain A) and positions 69, 71, 72 (chain B) move closer to the catalytic wall in the resistance ensemble in a symmetric fashion. Larger distances are at positions 10 (chain A) and 10, 21, 37, 54 (chain B). The elbow movement is not mirrored in chain A, however position 10 in chains A and B move away from the plane spanning the surface of the page, showing another way of active site expansion in addition to elbow flaring associated with the resistance ensemble. Residue 54 (chain B) is also seen to move away from the the binding cavity, but same does not occur under the replication *(SI Appendix*, Fig. S4). DRV’s resistance ensemble includes amongst other variations, the accessory DRMs 11I, 32I, 33F, 89V and the major DRMs 47V, 54LM, 84V.

For FPV (Fig.3C and 4C), smaller distances in the resistance ensemble are at positions 71 (chain A) and 69-72 (chain B), while larger distances in the resistance ensemble are at positions 15-17 (chain A) and 16, 73 (chain B). As in DRV lateral expansion is observed, but mainly involves the 10’s region in addition to the surface residue 73 in both replicates. The constriction behavior is also reproduced very closely in the replicate (*SI Appendix*, Fig. S4). The major DRM includes 84V while accessory DRMs include 10IVF, 32I, 46LI, 47V, 54LVM, 73S, 76V, 82TA, 90M in addition to other variations.

IDV (Fig.3D and 4D) displays smaller distances in its resistance ensemble at positions 63, 69-71 (chain A) and 71 (chain B). Larger distances are observed for the same ensemble at positions 16, 73 (chain A) and 16, 73, 93 (chain B). Upon replication, same residues were found to be involved in expansion, while only chain A showed the cantilever loop compaction towards the active site. Once more, the cantilever residue 73 is found to contribute to lateral widening in both chains. Replication identified identical residues involved in expansion at the 10’s loop from both chains, while those involved in compaction included residues 69-71 only from chain A (*SI Appendix*, S4). Major DRMs in the IDV resistance ensemble include 46LI, 82FTA, 84V and the accessory mutations 10IV, 20R, 32I, 36I, 54V, 71TV, 73SA, 76V, 77I, 90M.

Resistance in LPV (Fig.3E and 4E) was associated with smaller distances in the resistance ensemble at positions 70, 71 (chain A) and position 69-71 (chain B), while larger distances were located at positions 73, 93 (chain A) and positions 34, 36, 73 (chain B). Replicate runs are very concordant for residues involved in expansion with the exception of residues 36 and 81 in chain B, which rank differently despite displaying similar trends. Those involved in compaction again point to the cantilever residues of both chains, whereby residue 71 is replaced by 69 in the replicate. The major DRMs of the resistance ensemble consist of 32I, 47VA, 76V, 82SFTA while accessory DRMs comprise 10IFV, 20RM, 24I, 33F, 46LI, 50V, 53L, 54LTVMS, 63P, 71VT, 73S, 84V, 90M.

In the case of NFV (Fig.3F and 4F), shorter distances for the resistance ensemble were at positions 69-71 (chain A) and positions 70, 71 (chain B), while larger distances for the same ensemble were at positions 20, 36 (chain A) and at positions 20, 36, 73 (chain B). In the replicate run, a very similar profile is observed, however it would appear that residue 20 (part of the fulcrum) and 36 (close to elbow) are moving in concert during expansion. Contraction is observed as for other ARVs, close to the cantilever loop region. Major DRMs of the resistance ensemble consist of 30N, 90M and accessory DRMS consist of 10IF, 36I, 46LI, 71TV, 77I, 82FA, 84V, 88D.

SQV (Fig.3G and 4G) displays smaller distances in the resistance ensemble at positions 70, 71 (chain A) and positions 69, 71, 72 (chain B). Larger distances for the same ensemble are observed at positions 73, 89 (chain A) and positions 18, 20, 73 (chain B). Very similar symmetric compaction is observed at the fulcrum region as seen for other ARVs on both chains and the widening peak positions are also very similar despite a slightly changed degree ranking. DRMs for the resistance ensemble include the majors 48V, 90M and the accessory mutations 10I, 54LV, 62V, 71TV, 73S, 77I, 82A, 84V.

In the case of TPV (Fig.3H and 4H), smaller distances in the resistances are at positions 33, 60, 71 (chain A) and positions 70, 71 (chain B), while larger distances are at positions 16, 20 (chain A) and positions 15-17 (chain B). Replication reproduced compactions once more, close to the cantilever loop but also included the buried residue at position 33 on chain A, surrounded by the 80’s loop, the cantilever and the elbow regions. According to our simulation conditions, compaction at this region appears to be specific to TPV. Lateral expansions, though not identical, are also closely reproduced around the 10’s loop region. The resistance ensemble includes the major DRMs 47V, 58E, 74P, 82LT, 83D, 84V and the accessory DRMs 10V, 33F, 36IV, 43T, 46L, 54VM, 89VM.

We further investigate receptor backbone movement by comparing angle distributions occurring at protein *C_α_* atoms. Absolute conservation was observed at residue position 84 in only one of the replicates in Fig.5. Positions 75 and 84 however displayed strongly conserved larger angles in the resistance ensemble, including ATV, DRV, FPV, IDV, LPV, NFV and SQV. At 99% confidence), one-tailed t-tests did not detect any strong conservation of global angular behavior - both for the same drug replicated and across all drugs as shown by the non-reproducible clustering patterns in Figs.5 and 6. This supports the fact that the enzyme is very malleable, even in the closed conformation complexed with the drug and points to the fact that multiple residue arrangements along the backbone can lead to the same effect.

**Figure 5.**
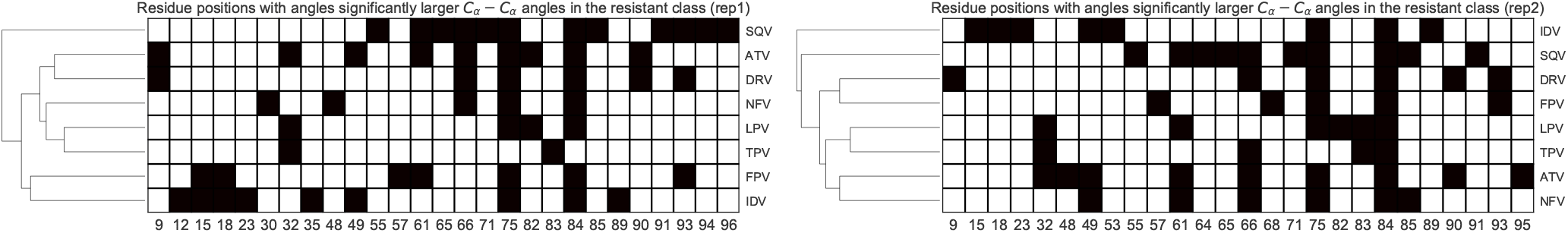
Heat map of residue positions with significantly larger *C_α_* angles in the resistant ensemble for each PI. The hierarchical cluster tree is displayed on the left. The first replicate is at the left and the second replicate is at the right.

**Figure 6.**
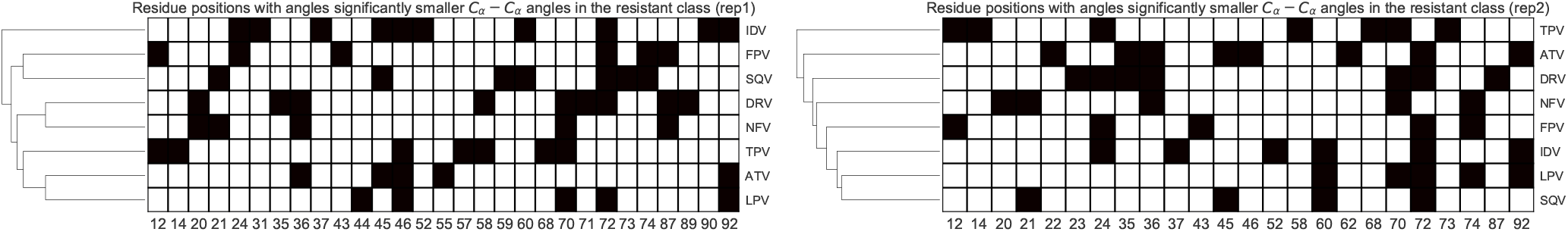
Heat map of residue positions with significantly smaller *C_α_* angles in the resistant ensemble for each PI. The hierarchical cluster tree is displayed on the left. The first replicate is at the left and the second replicate is at the right.

**In conclusion**, HIV protease inhibitors are used to delay the symptoms associated with late stages of the infection, however resistance is unrelenting due to the virus’s resilience to the current drug designs. Drug resistance patterns are complex. Nevertheless, our large scale simulations show that despite various DRMs and additional variations, lateral expansion and fulcrum compaction are conserved in the drug resistance state, both within and between different types of PIs. Analysis of the backbone motions hinted that there is no single angular trajectory leading to resistance, even for the same sequence. Knowledge of characteristic motions around similar energy wells may be an interesting and inexpensive route for supplementing extant drug resistance prediction approaches in HIV subtype B. Given phenotypically-labeled protease sequences from other subtypes, a similar experimental design may prove to be quite useful in improving prediction further, should similar conserved local motions prevail.

We have used MD and network centrality measures to identify common structural features in drug-resistant mutations of HIV protease. To our knowledge, this method is novel, although elastic network models were used to determine the functional effects of variants in other proteins^39^. We expect that our method will be useful in other cases for analyzing protein structural variations.

## Methods

### Dataset preparation

HIV subtype B protease sequence variants labeled with fold drug resistance ratios were obtained from the Stanford HIVdb unfiltered dataset^23^. These were reconstituted and filtered as explained in^24^. After ranking the sequences on the basis of decreasing average distance for each of the 8 PIs, 100 highly-resistant and 100 hyper-susceptible sequences were shortlisted, using cut-offs defined in^40^. These two classes of sequences are henceforth referred as to the resistant and the susceptible ensembles respectively. Sequences are provided in supplementary dataset *SI Appendix, dataset S5*. Pandas 0.21.0^41^ was used for dataset storage and manipulation. Seaborn 0.7.1 and matplotlib 2.1.0^42^ were used for plotting.

### Homology modelling

Modeller (version 9.16) was used to model each of the protein sequences in their closed conformation. High resolution (¡ 1.55 Å) crystal structure templates were retrieved for each of the 8 available PIs from the HIVdb dataset (PDB accessions: 3NU3, 3EL9, 2HS1, 2AVO, 2O4S, 3EL5, 2NMZ and 3SPK). Very slow refinement was used and model quality was assessed using z-DOPE scores.

### Ligand docking

Flexible ligand docking was performed using AutoDock Vina (version 1.1.2)^35^ to place each PI in its respective receptor variant using an exhaustiveness of 16. The docking center was picked from a saquinavir atom from template 2NMZ subsequently used as reference to align the totality of the homology models using ProDy^43^.

### Molecular dynamics

Receptors were protonated to pH7 using the PROPKA algorithm from PDB2PQR^44^, while parameters for the docked ligand poses were determined using ACPYPE^45^after full protonation using VEGA (version 3.1.1)^46^. All 8 x 200 complexes were prepared for molecular dynamics using GROMACS (version 2016.1)^47^. The AMBER03 forcefield was used with a short-range non-bonded interaction cut-off distance of 1.2nm. Long-range electrostatics were handled using the smooth Particle Mesh Ewald algorithm. Energy-minimization was performed using the method of steepest descent after neutralizing charges using 0.15M sodium chloride in SPC-modeled water within a triclinic periodic box. A 50ps temperature equilibration (at 310K) was followed by 50ps of pressure equilibration (1 atm) with time steps of 2fs and finally a 2ns production MD was performed at the same temperature, pressure and time step. All MD runs were distributed over a 2400-core queue with 24 cores per job using GNU Parallel (version 20160422)^48^, managed by the PBS Professional scheduler over the lengau cluster (Centre for High Performance Computing (CHPC)).

### Trajectory analysis

After generating MD trajectories, the proteins were centered and rotations/ translations were removed using the trjconv command in GROMACS. RMSD values were first evaluated to detect any failure in correcting periodic boundary conditions. These plots identified an initial period of fluctuation spanning the first 100ps, which were dropped from any subsequent analysis. *Rg* values were calculated to have an overview of the levels of compaction observed in the resistance ensemble compared to susceptible ensemble for each drug investigated. Thereafter, local analyses were performed: (a) Welsch t-tests were evaluated over pairwise residue distances across the ensembles. To do so, pairwise *Cβ* (and *C_α_* for glycine) atom distances from each trajectory were time-averaged within each ensemble. For each drug, each pairwise residue distance was aggregated into separate two-dimensional arrays - one for each ensemble. The t-tests were then performed between each analogous array at a 99% confidence level. (b) The angles between *C_α_* residues were computed for each complex within an ensemble and compared against the analogous array of angles in the other ensemble using t-tests. Only those angles corresponding to p-values above 2.5 standard deviations were retained for either of the larger or smaller angles in the resistance ensembles. Bonferroni correction was applied in both approaches to correct for multiple testing and reduce chances of false positives. Finally, the angles found to be significant for each drug were clustered by average linkage from the matrix of pairwise Euclidean distances. The MDTraj library (version 1.9.1)^49^ was used in Python 3.5 for trajectory distance and angle calculations. Numpy 1.13.3 and scipy 1.0.0^50^ were used for general computations and statistical tests respectively.

### Network analysis

Network graphs were built from nodes corresponding to *Cβ* (or glycine *C_α_*) atoms. Edges were obtained from significant p-values obtained from independent t-tests performed on arrays of time-averaged pairwise residue distances. In other words, each time-averaged pairwise distance 〈*D_ij_*〉 for a given protein concatenated to those of other proteins within the ensemble. Each array of 〈*D_ij_*〉 values is then compared to its corresponding position in the other ensemble of 〈*D^′^_ij_*〉 values using 2 sample t-tests. In order to expose more information, one-tailed tests were performed to determine whether distances are larger or smaller between the resistance ensembles. Same method was applied for all drugs. Finally the node degree centralities were calculated and the top 5 most central nodes for both higher and lower distances were shown as text labels for each drug. Network construction and analysis were performed using the NetworkX library (version 1.11)^51^. Edge mappings onto protein structures were generated using the the NGLview library (version 1.0)^52^.

## Acknowledgements

We thank the Centre for High Performance Computing (CHPC), South Africa for computational resources, and the National Research Foundation (NRF), South Africa for funding under grant 93690.

## Author contributions statement

Author contributions: O.S.A. designed and performed the research and wrote the first draft of the manuscript; Ö.T.B. and N.T.B. supervised the research and edited the manuscript.

